# A perceptual decision making EEG/fMRI data set

**DOI:** 10.1101/253047

**Authors:** Yasmin K. Georgie, Camillo Porcaro, Stephen D. Mayhew, Andrew P. Bagshaw, Dirk Ostwald

**Author notes:** Corresponding author: Dirk Ostwald.

## Abstract

We present a neuroimaging data set comprising behavioural, electroencephalographic (EEG), and functional magnetic resonance imaging (fMRI) data that were acquired from human subjects performing a perceptual decision making task. EEG data were acquired both independently and simultaneously with fMRI data. Potential data usages include the validation of biocomputational accounts of human perceptual decision making or the empirical validation of simultaneous EEG/fMRI data processing algorithms. The dataset is available from the Open Science Framework and organized according to the Brain Imaging Data Structure standard.

## Background & Summary

Perceptual decision making is the selection of one among a set of possible interpretations of a sensory event^1,2,3,4^. While some progress has been made in identifying the neural markers of human perceptual decisions over the last decade (for recent reviews, see Hanks & Summerfield ^5^ and Summerfield & Blangero^6^), a comprehensive biocomputational account of human perceptual decision making is still lacking. In this data descriptor, we report on a data set that may serve as an empirical testbed for such an account. The data set comprises behavioural, electroencephalographic (EEG), and functional magnetic resonance imaging (fMRI) data from a group of human subjects performing a visual perceptual decision making task. The task is similar to perceptual decision paradigms employed previously^7,8,9^ and involves the experimental manipulation of stimulus informativeness (coherence) and stimulus prioritization (spatial attention). Notably, EEG data were acquired both independently and simultaneously with fMRI data. The data reported herein thus allow for testing biocomputational accounts that are either constrained by each neuroimaging modality in isolation or by both modalities on a trial-by-trial basis^10,11,12^. Alternatively, the data set may also be viewed as an empirical validation data set for algorithmic data processing developments in simultaneous EEG/fMRI research, for which an artefact-free EEG standard is desired^13,14,15^. We have previously explored these viewpoints using the current data in Ostwald *et al.*^16^ from a data-driven, information theoretic perspective^17,18^. Below, we report on conventional response time and accuracy, event-related potential, and GLM-based fMRI data analyses of the data set. These technical validation analyses indicate the presence of both standard neural markers of perceptual decisions, as well as of paradigm-specific effects. To make the data set accessible for reuse, it is organized according to the Brain Imaging Data Structure (BIDS), a recently developed standard for neuroimaging data sharing^19^ (http://bids.neuroimaging.io). The data set is made available via the Open Science Framework^20^ (https://osf.io).

## Overview

Behavioural, EEG, and fMRI data were acquired from a cohort of subjects performing a visual perceptual decision making task.The task was embedded in a 2×2 factorial within-subject de-sign with factors visual stimulus coherence and spatial prioritization (attention) as detailed below (Figure 1a and b). Subjects performed the task in two separate experiments. Specifically, after preparation of the EEG cap, subjects first performed two runs of the task outside the MR scanner while behavioural and EEG data were recorded. We refer to the data of this experiment as “EEG only” and “outside MRT” data. After a short break, subjects were situated in the MR scanner and, simultaneously with the acquisition of behavioural and fMRI data, EEG data was acquired in five runs of the task. We refer to the data of this experiment as “simultaneous EEG/fMRI” and “inside MRT” data. Table 1 provides an overview of the components of the data set that could successfully be recorded from each subject.

**Figure 1.**
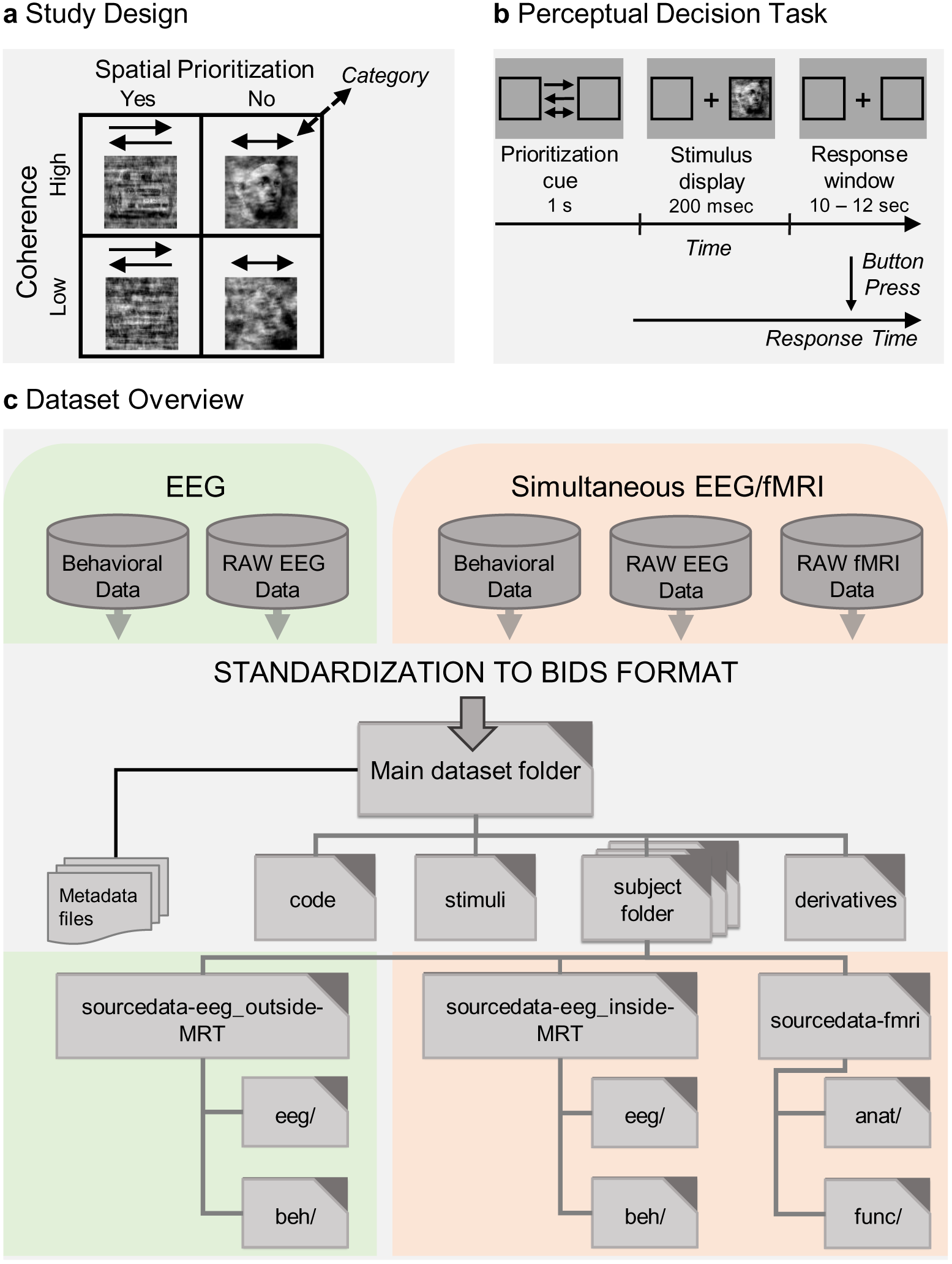
Study design, perceptual decision task, and dataset overview. **a.** 2x2 factorial experimental design with factors stimulus coherence (low, high) and spatial prioritization (yes, no). On each trial of the experiment, the subject was presented with a face or car stimulus, which had been manipulated according to its spatial phase coherence. Simultaneously, the subject was prompted to either spatially prioritize the stimulus display or not. The stimulus category (face or car) that the subject was asked to discriminate, was manipulated orthogonally to the other factors, b. Single experimental trial. Prior to the presentation of the stimulus, a one-headed arrow could indicate the hemifield of the subsequent stimulus presentation for spatial prioritization. Alternatively, a two-headed arrow was uninformative with respect to the location of the upcoming stimulus in the no spatial prioritization condition. The cueing arrow was shown continuously for 1 s pre-stimulus, the stimulus itself was shown for 200 ms. The subject was asked to respond as quickly and as accurately as possible with no restrictions on the response window. The inter-trial interval was 0-300 ms for the EEG only and 10-12 s for the simultaneous EEG/fMPJ recordings, c. Schematic overview of the study and dataset. EEG and simultaneous EEG/fMPJ data were acquired over two data acquisition sessions. The resulting imaging and behavioural data were standardized into BIDS format. The figure depicts the folder structure as available from the Open Science Framework.

**Table 1.** Subject data set inventory. Green ticks represent successfully recorded data set components, blue crosses represent absent data set components. Overall, the dataset includes behavioural data acquired simultaneously with EEG data outside of the MR environment from 16 subjects (BEH outside), behavioural data acquired simultaneously with EEG and fMRI data from 16 subjects (BEH inside), EEG data recorded outside the MR environment from 16 subjects (EEG outside), EEG data recorded simultaneously with fMRI data from 17 subjects (EEG inside), and fMRI data from 16 subjects (fMRI).

| Subject | BEH outside | BEH inside | EEG outside | EEG inside | fMRI |
| --- | --- | --- | --- | --- | --- |
| sub-001 | ✓ | ✗ | ✓ | ✓ | ✗ |
| sub-002 | ✗ | ✓ | ✗ | ✓ | ✓ |
| sub-003 | ✓ | ✓ | ✓ | ✓ | ✓ |
| sub-004 | ✓ | ✓ | ✓ | ✓ | ✓ |
| sub-005 | ✓ | ✓ | ✓ | ✓ | ✓ |
| sub-006 | ✓ | ✓ | ✓ | ✓ | ✓ |
| sub-007 | ✓ | ✓ | ✓ | ✓ | ✓ |
| sub-008 | ✓ | ✓ | ✓ | ✓ | ✓ |
| sub-009 | ✓ | ✓ | ✓ | ✓ | ✓ |
| sub-010 | ✓ | ✓ | ✓ | ✓ | ✓ |
| sub-011 | ✓ | ✓ | ✓ | ✓ | ✓ |
| sub-012 | ✓ | ✓ | ✓ | ✓ | ✓ |
| sub-013 | ✓ | ✓ | ✓ | ✓ | ✓ |
| sub-014 | ✓ | ✓ | ✓ | ✓ | ✓ |
| sub-015 | ✓ | ✓ | ✓ | ✓ | ✓ |
| sub-016 | ✓ | ✓ | ✓ | ✓ | ✓ |
| sub-017 | ✓ | ✓ | ✓ | ✓ | ✓ |

## Participants

Seventeen participants (8 female, mean age 25.9 years, range 20-33 years, 2 left-handed) were recruited from the University of Birmingham campus. Participants were paid 7.5 £ per hour and their travel expenses were reimbursed. All participants had normal or corrected to normal vision, no history of neurological disorders and provided written informed consent. The study was approved by the Science, Technology, Engineering and Mathematics Ethical Review Committee of the University of Birmingham.

## Stimuli

### Original stimulus set

The original stimulus set consisted of 18 pictures of cars and 18 pictures of faces, similar to the stimulus set used in^7,8,9^. The original car images were obtained from the website of the Laboratory for Intelligent Imaging and Neural Computing (Prof. Paul Sajda, http://liinc.bme.Columbia.edu). The original face images were obtained from the Max Planck face database^21^ (http://faces.kyb.tuebingen.mpg.de). The two original image sets were matched for the number of frontal, left lateral, and right lateral car and face views, respectively. All images were converted from their native format to bitmap format (.bmp) and the resulting 256 × 256 matrices were saved with 8 bit depth. The two original stimulus sets were matched for their mean driving luminance and contrast as assessed by a one-way ANOVA with factor “image category” and levels “face” and “car” (mean driving luminance: *F*_(1,34)_ = 0.08, *p* = 0.78, contrast: *F*_(1,34)_ = 0.23, *p* = 0.64).

### Phase-scrambled stimulus set

To manipulate the informativeness of the images, the images’ spatial phase spectra were linearly weighted with a phase spectrum of a uniform noise image using the weighted mean phase technique described in Dakin *et al.*^22^. With the original phase spectrum of an image given by *ϕ_0_*, the scrambled phase spectrum *ϕ_s_* was computed as

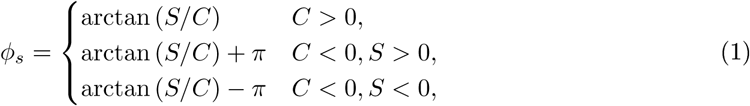

where

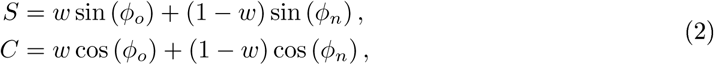

*ϕ_n_* is the phase spectrum of a random noise image, created by sampling each pixel’s value uniformly from the interval [0,1] using Matlab’s *rand.m* function, and *w* ∈ [0,1] is the signal-to-noise weighting coefficient. Stimuli with weighting coefficients *w_1_* = 0.9 (high coherence) and *w_2_* = 0.5 (low coherence) were chosen for the experiment in order to elicit reliable differences in the response times for either stimulus class, while still allowing accurate performance of the task. The stimulus set generation is implemented in the Matlab (The MathWorks, NA) function *pdm_ experiment _ stimulus _ creation.m,* which is available with the dataset along with all custom generated programming code.

### Experimental procedure

Subjects performed a perceptual decision task in a 2 × 2 factorial within-subject design with experimental factors “stimulus coherence” (with levels “low” and “high”) and “spatial prioritization” (with level “yes” and “no”) (Figure 1a). On each trial, a visual stimulus depicting either a face or a car was presented in one visual hemifield with a left/right eccentricity of the stimulus center of 11 degrees of visual angle and a stimulus extension of 9 degrees of visual angle. Individual stimuli were presented for 200 ms and the subject was asked to indicate via a button press whether the stimulus depicted a face or a car. For the button presses, subjects used their right index and middle finger for the two categories, and the mapping from stimulus category to response button was counterbalanced across subjects. As described above, the informativeness of the visual stimulus was manipulated by altering the phase coherence of its spatial frequency spectrum resulting in low and high stimulus coherence trials. On half of the trials, a cueing arrow shown continuously for 1 s prior to the stimulus indicated in which visual hemifield the stimulus would be presented (Figure 1b). Subjects were asked to allocate their spatial attention to the respective visual hemifield, while maintaining steady central fixation (spatial prioritization condition). On the other half of the trials, the two-headed cuing arrow was uninformative and the stimulus was presented randomly in either visual hemifield (no spatial prioritization condition). We refer to the attention factor as “spatial prioritization” rather than “spatial attention”, because in the no spatial prioritization condition, spatial attention is equally directed to both potential stimulus displays. This differs from classical spatial attention experiments, in which evoked responses for spatially attended stimuli are typically contrasted with spatially inhibited stimuli^23^. Face and car stimuli were equally distributed across the four experimental conditions. The stimulus presentation order was randomized. Subjects were asked to respond as quickly and as accurately as possible with an emphasis on responding as quickly as possible and to maintain stable fixation on the central fixation cross throughout the experiment. For the EEG recordings outside of the scanner, data from 72 trials for each of the four conditions (half of them face stimuli) were recorded with an inter-trial interval randomized between 0 ms and 300 ms. Here, the data acquisition was split into two experimental runs of approximately 10 minutes each. For the simultaneous EEG/fMRI recordings data from 90 trials for each of the four conditions (half of them face stimuli) were recorded with an inter-trial interval discretely randomized between 10 and 12 s. This long inter-trial interval was chosen to obtain reliable recordings of single-trial haemodynamic responses. The 90 trials per condition were split into five experimental runs, each lasting approximately 14 minutes. The EEG only and EEG/fMRI experiments were presented under Matlab using Psychtoolbox-3 (http://psychtoolbox.org) with the functions *pdm_experiment_eeg_only.m* and *pdm_experiment_eeg_fmri.m,* respectively.

### EEG data acquisition

EEG data were recorded using a 64-channel MR-compatible EEG system (Brain Amp MR Plus, Brain Products, Munich, Germany), which incorporates current limiting resistors of 5 *k*Ω at the amplifier input and in each electrode. The EEG cap consisted of 62 scalp electrodes distributed according to the 10-20 system^24^ and two additional electrodes, one of which was attached approximately 2 cm below the left collarbone for recording the ECG, the other one of which was attached below the left eye (on the lower orbital portion of the orbicularis oculi muscle) for detection of eyeblink artefacts. Data were sampled at 5000 Hz. Impedance at all recording electrodes was less than 20 kll For simultaneous EEG/fMRI recordings, the EEG data acquisition setup clock was synchronized with the MRI scanner clock using Brain Products SyncBox, resulting in exactly 10,000 data points per EPI-TR interval. The EEG setup was identical for the recordings outside and inside the MR scanner.

### MRI data acquisition

The simultaneous EEG-fMRI experiment was conducted at the Birmingham University Imaging Centre using a 3T Philips Achieva MRI scanner. An initial T1-weighted anatomical scan (1 mm isotropic voxels) and T2*-weighted functional data were collected with an eight-channel phased-array SENSE head coil. EPI data (gradient echo-pulse sequence) were acquired from 32 slices (3x3x4 mm resolution, TR 2,000 ms, TE 35 ms, SENSE factor 2, flip angle 80 deg). Slices were oriented parallel to the AC-PC axis of the subject’s brain and positioned to cover the entire brain space.

## Data Records

The data set is available from the Open Science Framework via the private project https://osf.io/q4t8k/. It is organized according to the Brain Imaging Data Structure (BIDS) specification^19^. Extensive documentation of this neuroimaging data standard, including its metadata specifications for behavioural, EEG, and fMRI data, is available from the BIDS website (http://bids.neuroimaging.io). Table 2 provides an overview of the data set organization. We here limit the description of the data records to the top level in the first column. At this level, the data set contains metadata files, a *code/* folder hosting the custom-written Matlab code for the study and the technical validation analyses reported herein, and a *stimuli/* folder hosting the visual stimulus set. The behavioural and neuroimaging data itself is organized in a subject-wise manner in the *sub<subject id>/* folders, and substructured into further folders as documented in Table 2. Finally, the folder *derivatives/* contains EEG data files from inside the scanner that underwent run-wise segmentation and gradient and ballistocardiogram artefact removal.

**Table 2.**
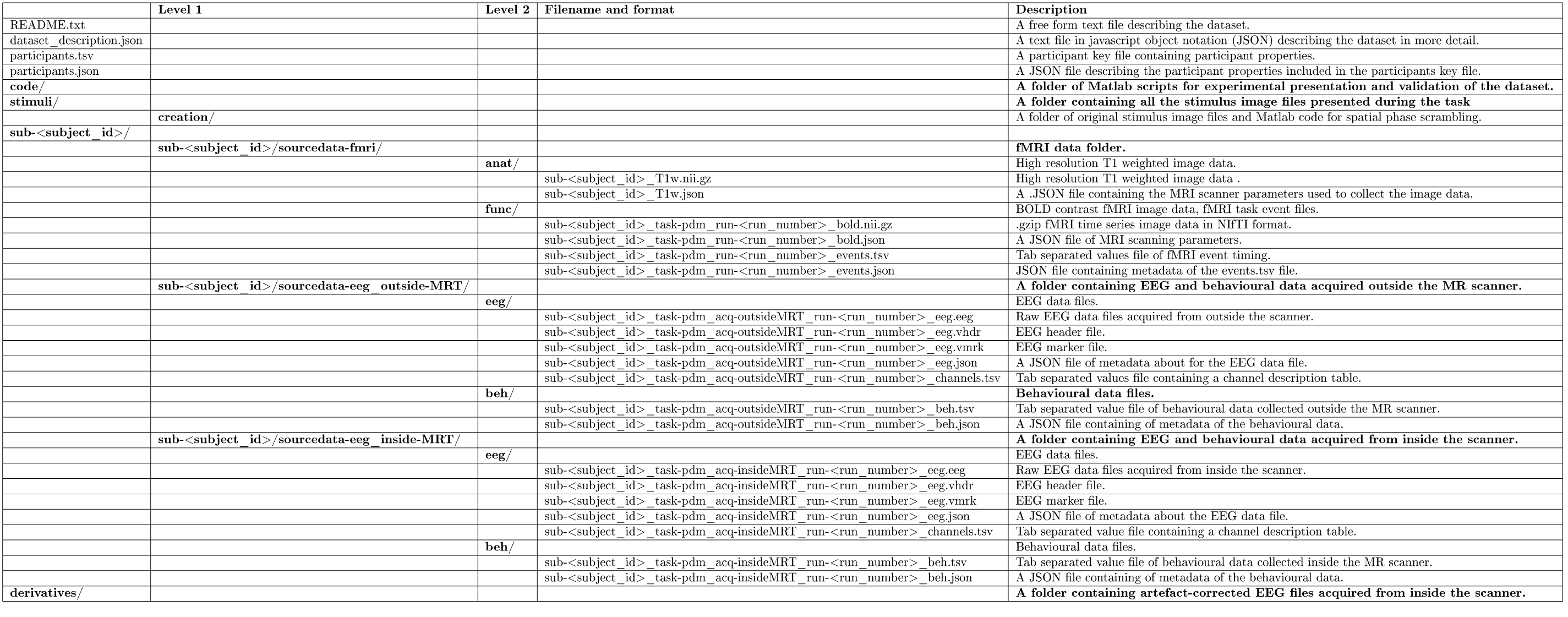
Data records overview. The table depicts and documents the data set structure as available from the Open Science Framework. The data organization is based on the Brain Imaging Data Structure (BIDS), extensive documentation of which is available from http://bids.neuroimaging.io.

## Technical validation

### Behavioural data

To validate the behavioural data quality, response times and response accuracy were evaluated for both the EEG only and EEG/fMRI experiment (Figures 2a and b). In both experiments, faster median response times were observed for the high stimulus coherence compared to the low stimulus coherence condition and for the spatial prioritization compared to the no spatial prioritization condition (Figure 2a). Equivalently, response accuracy increased with stimulus coherence and spatial prioritization (Figure 2b). The observed behavioural pattern was identical between the EEG and EEG/fMRI experiments. However, the MR scanner environment led to an overall increase in response times and a decrease in performance accuracy (mean response time across conditions: 403 ms (±20 standard error of the mean) for the EEG experiment vs. 727 ms (±33) for the EEG/fMRI experiment, accuracy across conditions: 91 (±1)% correct for the EEG experiment vs. 85 (±1)% correct for the EEG/fMRI experiment). The reliability of the experimental manipulation on the behavioural responses was assessed with a two-way repeated-measures ANOVA with factors stimulus coherence and spatial prioritization. A level of *p <* 0.005 was used for labelling effects “statistically significant”^25,26^. For response times of correct response trials, this ANOVA revealed significant main effects of stimulus coherence (EEG: *F*_(1,15)_ = 41.1, *p <* 0.001, EEG/fMRI: *F*_(1,15)_ = 23.3,*p <* 0.001) and spatial prioritization (EEG: *F_(1,15)_* = 14.1,*p* = 0.002, EEG/fMRI: *F*_(1,15)_ = 29.3,*p <* 0.001). No significant interactions were observed (EEG: *F*_(1,15)_ = 0.15,*p* = 0.71, EEG/fMRI: *F*_(1,15)_ = 0.86,*p =* 0.37). For accuracy, a significant main effect of stimulus coherence (EEG: *F*_(1,15)_ =52.5,*p <* 0.001, EEG/fMRI: F_(1,15)_ = 128.5,*p <* 0.001) and statistically suggestive effects of spatial prioritization (EEG: *F*_(1,l5)_ = 4.37,*p* = 0.05, EEG/fMRI: *F*_(1,15)_ = 4.1,*p* = 0.06) were observed. Like for response times, there were no significant interactions (EEG: *F*_(1,15)_ = 0.02, *p =* 0.89, EEG/fMRI: *F*_(1,15)_ = 1.6,*p =* 0.22). In summary, the behavioural data from both experiments exhibit commonly observed effects of stimulus coherence and spatial prioritization in perceptual decision making tasks^23,6,27^. The behavioural analyses reported here are implemented in the function *pdm_behaviour_analysis.m*.

**Figure 2.**
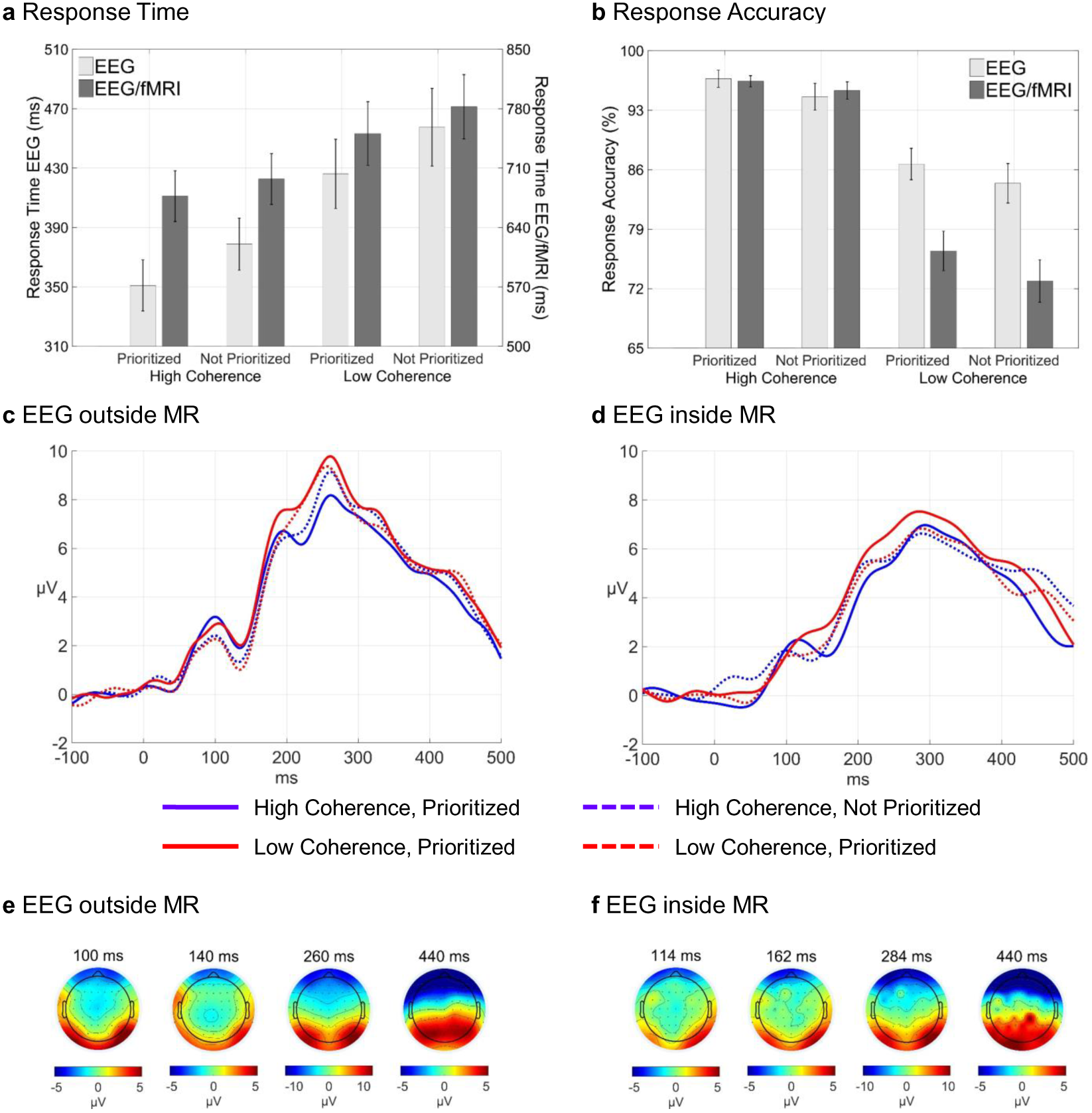
Behavioural and EEG data validation. **a.** Response time. Bars depict the average median response times across subjects, error bars ± standard error of the mean (SEM). Light grey bars depict behavioural data from the EEG data set recorded outside the MR scanner (EEG), dark grey bars depict behavioural data from the simultaneous EEG/fMRI recordings, **b.** Response accuracy. Bars depict the response accuracy in percent correct across subjects, error bars ± standard error of the mean (SEM). As for panel a, light grey bars depict behavioural data from the EEG data set recorded outside the MR scanner (EEG), dark grey bars depict behavioural data from the simultaneous EEG/fMRI recordings, **c** and **d.** Condition-specific group average ERPs for contralateral trials, pooled over electrodes 02, P04, P08 for left hemifield trials and Ol, P03, P07, for right hemifield trials for the EEG data recorded outside the MR scanner (c) and simultaneously with the fMRI data (d). **e** and **f.** Topography plots of the ERP data averaged over conditions for four selected post-stimulus time points for both EEG data sets. The main positive deflections at all time points were observed in the set of parieto-occipital electrodes (02, Ol, PQ8, PQ7, PQ4, PQ3) evaluated in the ERP analyses.

### EEG data

To validate the EEG data quality in both experiments, event-related potentials (ERPs) were computed for all conditions (Figure 2c-f). Before event-related averaging, data from the EEG recordings outside the MR scanner were down-sampled to 500 Hz, re-referenced to the average of all electrodes, and band-pass filtered from 0.5 to 25 Hz using SPM12 (V6906) (http://www.fil.ion.ucl.ac.uk/spm). To correct for MR-induced artefacts, EEG data from the simultaneous EEG/fMRI recordings were preprocessed using Brain Vision Analyzer 2.0. Specifically, after segmentation into the five data acquisition sessions, MR gradient and ballistocardiogram artefacts were removed and the data were down-sampled to 500 Hz. Subsequently, the data were imported to SPM12 (V6906), re-referenced to the average of all electrodes, and band-pass filtered from 0.53 to 25 Hz. After these preprocessing steps, the data from both experiments were epoched with a peri-stimulus time window of −100 ms to 500 ms and baseline corrected. The stimulus presentation onset times for the combination of left and right visual hemifield trials with the four experimental conditions were selected as events of interest. Upon data epoching and averaging over trials, this selection yielded a set of eight ERPs per subject and electrode. Figure 2c depicts the condition-specific group average ERPs averaged over electrodes 02, P04, and P08 for left visual hemifield trials and electrodes 01, P03, and P07 for right hemifield trials for the EEG data recorded outside the MR scanner. For all conditions, clear early (100 ms, P100) and late (270 ms, P300) positive potential deflections, separated by a negative potential deflection (140 ms, N140), could be identified. Obvious condition-specific effects included an increase of the P100 amplitude with spatial prioritization and an increase of the P300 amplitude with a decrease of stimulus coherence. Similarly, Figure 2d depicts the condition-specific group average ERPs averaged over electrodes 02, P04, and P08 for left visual hemifield trials and electrodes 01, P03, and P07 for right hemifield trials for the EEG data recorded simultaneously with the fMRI data. As for the EEG only experiment, condition-specific effects include an increase of the P100 amplitude with spatial prioritization and an increase of the P300 amplitude with a decrease of stimulus coherence.

Overall, the amplitude variations in the EEG/fMRI data are less pronounced and slightly delayed when compared to the EEG only data. Finally, to asses the topographic expression of the event-related responses, the ERP data were averaged over experimental conditions and projected onto a scalp representation at selected time-points using Fieldtrip’s *topoplot.m* function^28^. Here, for both data sets the strongest positive deflections for the time points of interest were observed for the set of posterior parieto-occipital electrodes selected for the reported ERPs (Figures 2e and 2f). In summary, the EEG data from both recordings yield standard ERP waveforms for visuomotor reaction tasks and reproduce known effects of stimulus coherence and spatial prioritization ^23,27,29^. The EEG data analyses reported here are implemented in the functions *pdm_ erp_ analysis_ eeg_ only.m* and *pdm_ erp_ analysis_ eeg_fmri.m.*

### fMRI data

To validate the fMRI data quality, a mass-univariate summary-statistics GLM analysis was performed that assessed condition-induced effects at the group-level. SPM12 (V6906) was used for both fMRI data preprocessing and statistical modelling. Prior to GLM parameter estimation at the subject-level, fMRI data were motion-corrected by realigning EPI volumes to the first volume of the first run of a given subject, normalized to MNI spaced using the SPM MNI-EPI template, re-interpolated to 2 mm isotropic voxel size, and smoothed using an 8 mm isotropic Gaussian kernel. The first-level GLM design matrix for each subject was then specified in run-wise, block-diagonal form. Here, each block comprised the four condition-specific stimulus onset functions, convolved with the canonical haemodynamic response function, in the column-wise order: high stimulus coherence/spatial prioritization, high stimulus coherence/no spatial prioritization, low stimulus coherence/spatial prioritization, low stimulus coherence/no spatial prioritization (Figure 3a). Per SPM defaults, the design matrices additionally comprised a constant run offset and a cosine basis function set implementing a temporal high-pass filter with a cut-off of frequency 1/128 Hz. High-frequency residual error correlations were accounted for by SPM’s default of approximating a first-order autoregressive process with white noise using parameterized covariance basis functions^30^. GLM beta and covariance component parameters were then estimated using SPM’s restricted maximum likelihood estimation scheme^31,32^. Beta parameter estimate contrast images were evaluated for the directed main effects “All Stimuli > Baseline” (contrast weight vector [111 1], reproduced over sessions), “High Coherence > Low Coherence” (contrast weight vector [11-1 −1]), “Low Coherence > High Coherence” (contrast weight vector [−1 −1 1 1]), “Spatial Prioritization > No Spatial Prioritization” (contrast weight vector [1 −1 1 −1]), and “No Spatial Prioritization > Spatial Prioritization” (contrast weight vector [−1 1 −1 1]). Finally, the resulting beta parameter contrast images evaluated at the group-level using one-sample t-tests.

Figures 3b-e visualize the results of the group-level GLM analysis. As expected, contrasting all stimulus onsets across conditions with the GLM baseline resulted in widespread cortical and subcortical activation, including activity clusters in the fusiform gyrus, the superior parietal lobule, the left precentral gyrus, the insula, the inferior frontal gyrus, the anterior cingulate cortex, and the thalamus (Figure 3b). Given the general nature of this “task vs. no task” contrast, we do not report futher statistics for this comparison. For the directed main effects of stimulus coherence, testing for higher BOLD activity in high coherence compared to low coherence stimulus trials yielded statistically significant activation clusters at a family-wise error corrected level of p < 0.005 in the left superior frontal and middle gyri, as well as in the left precuneus (Figure 3c, Table 3). Testing for higher BOLD activity for the reverse direction of low coherence as compared to high coherence stimuli yielded statistically significant activation clusters in the right anterior cingulate gyrus and the right inferior frontal gyrus. Activations in the superior parietal lobule, as well as the left and right frontal eye-fields were marginally statistically suggestive (Figure 3d, Table 3). For the directed main effects of spatial prioritization, a cluster in the left superior parietal lobule survived a cluster-defining threshold of p < 0.001. However, this cluster was not significant at the family-wise error corrected cluster level (Figure 3e, Table 3). Finally, no activity clusters were detected for the reverse main effect of no spatial prioritization as compared to spatial prioritization trials at a cluster defining threshold of p < 0.001. In summary, evaluation of the fMRI data resulted in activations of areas known to be involved in the processing of visual perceptual decisions ^6,4,33^. The fMRI data analyses reported here are implemented in the functions *pdm_fmri_preprocessing.m, pdm_fmri_glm_first_level.m,* and *pdm_fmri_glm_second_level.m.*

**Figure 3.**
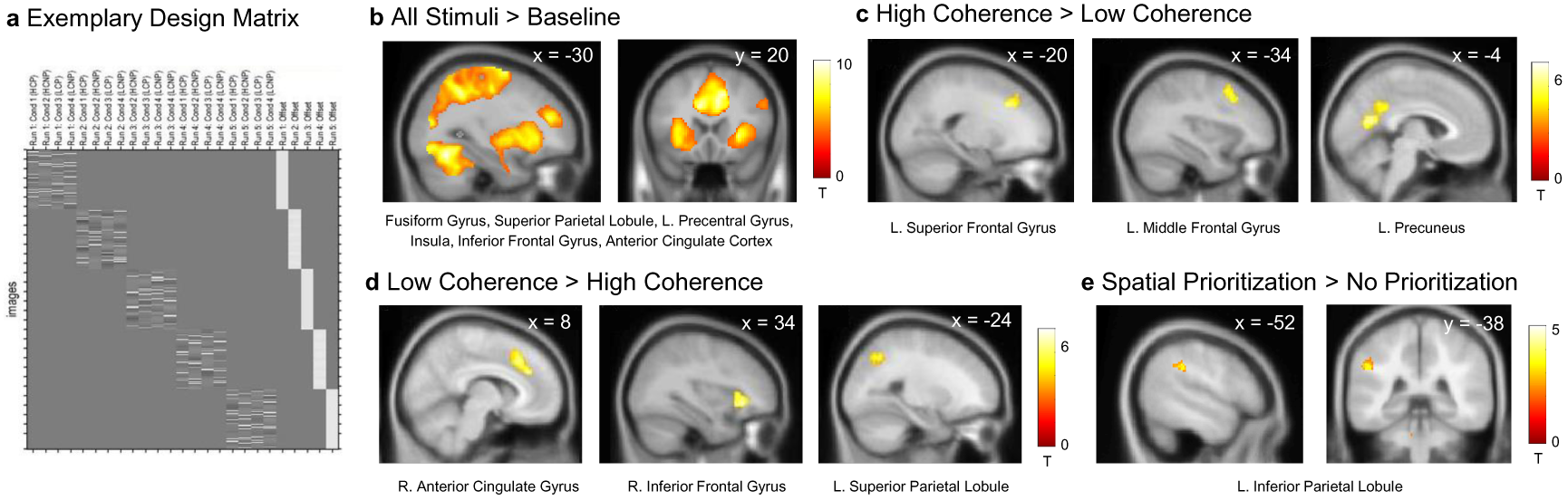
fMRI data validation. **a.** Exemplary first-level design matrix of a single subject. The design matrix comprised 20 regressors as columns corresponding to the four experimental conditions for each of the five experimental runs in the order high stimulus coherence/prioritized (HCP), high stimulus coherence/not prioritized (HCNP), low stimulus coherence/prioritized (LCP), and low stimulus coherence/not prioritized (LCNP) and five columns of constant run offsets. The design matrix’ 2,025 rows correspond to the 405 MPJ images acquired in each of the five experimental runs. **b-e.** Visualization of BOLD activation clusters for the directed main effects of the experiment. T-value statistical parametric maps are overlaid on the SPM average Tf image. All T-value maps are thresholded at a cluster-defining threshold of p < 0.001. The upper right corner insets denote the MNI coordinate of the respective slice. Anatomical labels are based on the WFU PickAtlas^34^.

**Table 3.**
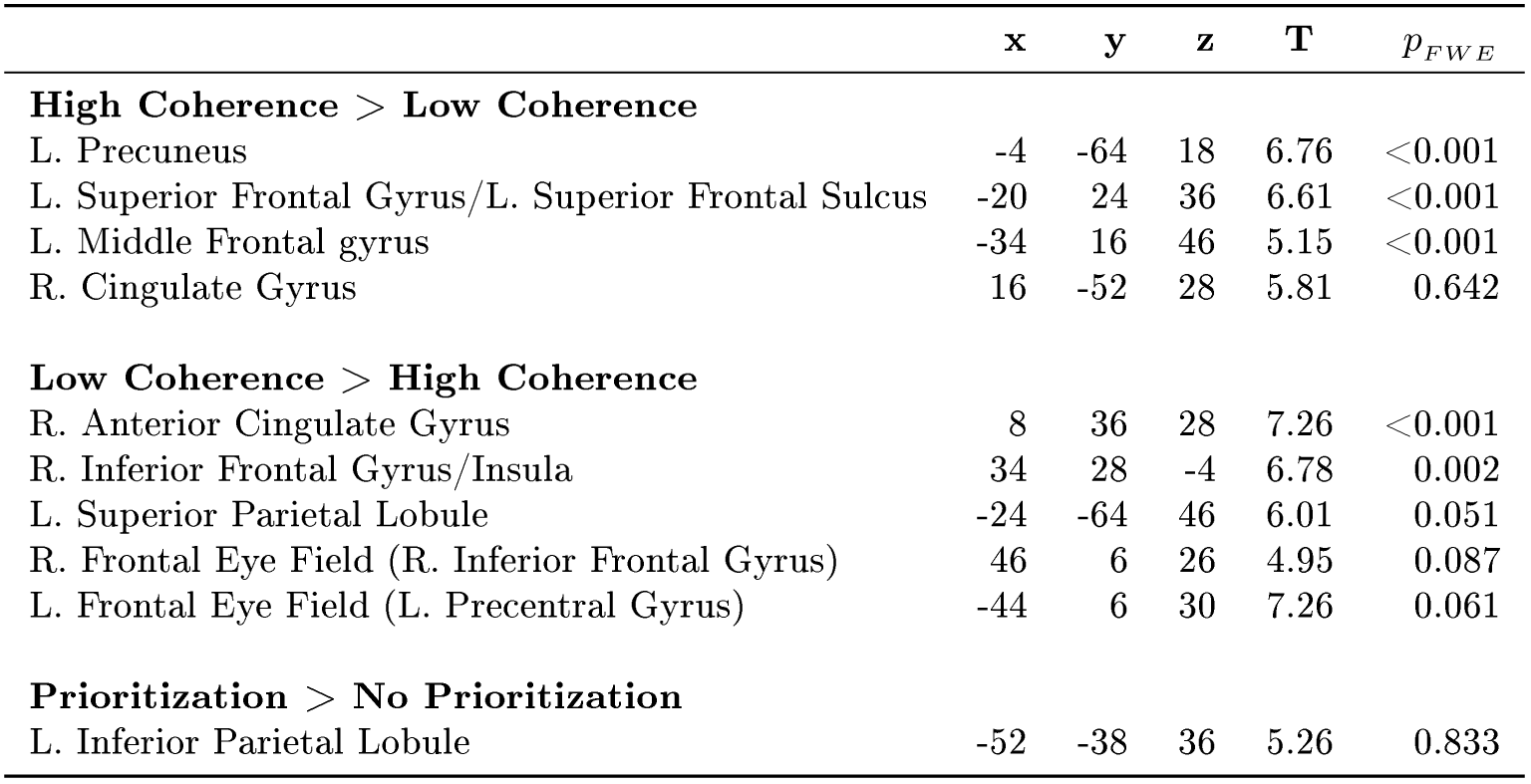
Experimental main effects group-level fMRI results. The table lists a cluster’s anatomical label according to the WFU PickAtlas^34^ atlas, its center of gravity in MNI coordinates, the T-value of its peak voxel, and the cluster level family-wise corrected p-value based on a cluster-defining threshold of p < 0.001 with an cluster extent threshold of 0 voxels^35^.

### Conclusion

In summary, the data set presented here provides a comprehensive representation of the neural processes underlying perceptual decision making as afforded by non-invasive neuroimaging techniques. Basic technical validation analyses suggest the presence of commonly observed experimental effects of stimulus coherence and spatial prioritization at the behavioural, EEG, and fMRI data level. The data set may thus be suited for both the development and validation of novel data analytical techniques, as well as for providing new insights into the neural mechanisms of perceptual decision making.

## Usage Notes

Because the data were donated by human participants, ethical considerations require some limitations on the access and reuse of the data.To obtain data access, potential data users should contact the corresponding author with a brief statement to which purpose the data will be used. Upon positive review of their request, data users are required to sign a data use agreement, which is available from https://osf.io/hkevu. Data users will then be added as collaborator to the Open Science Framework project hosting the data set and can download the data. We kindly request data users to follow the ODC Attribution/Share-Alike Community Norms (https://opendatacommons.org/norms/odc-by-sa), and acknowledge Yasmin K. Georgie, Camillo Porcaro, Stephen D. Mayhew, Andrew P. Bagshaw, and Dirk Ostwald in any publication derived from these data, citing this paper and Data Citation 1 as the source of the data.

## Acknowledgements

This work was supported by grant number EP/F023057/1 from the UK Engineering and Physical Sciences Research Council (EPSRC) and the Dr. Hadwen Trust.

## Author Contributions

Conceived and designed the experiments: D.O. and A.P.B.; acquired the data: D.O., S.M., C.P., and A.P.B.; contributed analysis tools: C.P.; analysed the data and wrote the paper: Y.K.G., D.O.

## References

1. W. T. Newsome, K. H. Britten, and J. A. Movshon, “Neuronal correlates of a perceptual decision,” Nature, vol. 341, no. 6237, pp. 52–54, 1989.

2. J. I. Gold and M. N. Shadlen, “The neural basis of decision making,” Annu. Rev. Neurosci., vol. 30, pp. 535–574, 2007.

3. H. R. Heekeren, S. Marrett, and L. G. Ungerleider, “The neural systems that mediate human perceptual decision making,” Nature Reviews Neuroscience, vol. 9, no. 6, pp. 467–479, 2008.

4. M. Mulder, L. Van Maanen, and B. Forstmann, “Perceptual decision neurosciences-a model-based review,” Neuroscience, vol. 277, pp. 872–884, 2014.

5. T. D. Hanks and C. Summerfield, “Perceptual decision making in rodents, monkeys, and humans,” Neuron, vol. 93, no. 1, pp. 15–31, 2017.

6. B. A. Summerfield C., Decision Neuroscience: An Integrative Perspective, Perceptual Decision-Making: What Do We Know, and What Do We Not Know?, pp. 149 – 162. Academic Press, 2017.

7. M. G. Philiastides, R. Ratcliff, and P. Sajda, “Neural representation of task difficulty and decision making during perceptual categorization: a timing diagram,” Journal of Neuroscience, vol. 26, no. 35, pp. 8965–8975, 2006.

8. M. G. Philiastides and P. Sajda, “Temporal characterization of the neural correlates of perceptual decision making in the human brain,” Cerebral Cortex, vol. 16, no. 4, pp. 509–518, 2006.

9. M. G. Philiastides and P. Sajda, “EEG-informed fMRI reveals spatiotemporal characteristics of perceptual decision making,” Journal of Neuroscience, vol. 27, no. 48, pp. 13082–13091, 2007.

10. J. Jorge, W. Van Der Zwaag, and P. Figueiredo, “EEG-fMRI integration for the study of human brain function,” Neurolmage, vol. 102, pp. 24–34, 2014.

11. M. Rosa, J. Daunizeau, and K. J. Friston, “EEG-fMRI integration: a critical review of biophysical modeling and data analysis approaches,” Journal of Integrative Neuroscience, vol. 9, no. 04, pp. 453–476, 2010.

12. S. Debener, M. Ullsperger, M. Siegel, and A. K. Engel, “Single-trial EEG-fMRI reveals the dynamics of cognitive function,” Trends in cognitive sciences, vol. 10, no. 12, pp. 558–563, 2006.

13. M. Ullsperger and S. Debener, Simultaneous EEG and fMRI: recording, analysis, and application. Oxford University Press, 2010.

14. C. Mulert and L. Lemieux, EEG-fMRI: physiological basis, technique, and applications. Springer Science & Business Media, 2009.

15. A. P. Bagshaw and T. Warbrick, “Single trial variability of eeg and fmri responses to visual stimuli,” Neurolmage, vol. 38, no. 2, pp. 280–292, 2007.

16. D. Ostwald, C. Porcaro, S. D. Mayhew, and A. P. Bagshaw, “EEG-fMRI based information theoretic characterization of the human perceptual decision system,” PLoS ONE, vol. 7, no. 4, p. e33896, 2012.

17. D. Ostwald, C. Porcaro, and A. P. Bagshaw, “An information theoretic approach to EEG-fMRI integration of visually evoked responses,” Neurolmage, vol. 49, no. 1, pp. 498–516, 2010.

18. D. Ostwald and A. P. Bagshaw, “Information theoretic approaches to functional neuroimaging,” Magnetic resonance imaging, vol. 29, no. 10, pp. 1417–1428, 2011.

19. K. J. Gorgolewski, T. Auer, V. D. Calhoun, R. C. Craddock, S. Das, E. P. Duff, G. Flandin, S. S. Ghosh, T. Glatard, Y. O. Halchenko, et al., “The brain imaging data structure, a format for organizing and describing outputs of neuroimaging experiments,” Scientific Data, vol. 3, p. 160044, 2016.

20. B. A. Nosek, G. Alter, G. C. Banks, D. Borsboom, S. D. Bowman, S. J. Breckler, S. Buck, C. D. Chambers, G. Chin, G. Christensen, et al., “Promoting an open research culture,” Science, vol. 348, no. 6242, pp. 1422–1425, 2015.

21. N. F. Troje and H. H. Bülthoff, “Face recognition under varying poses: The role of texture and shape,” Vision research, vol. 36, no. 12, pp. 1761–1771, 1996.

22. S. Dakin, R. Hess, T. Ledgeway, and R. Achtman, “What causes non-monotonic tuning of fMRI response to noisy images?,” Current Biology, vol. 12, no. 14, 2002.

23. S. Luck and S. Hillyard, The New Cognitive Neurosciences, The Operation of Selective Attention at Multiple Stages of Processing: Evidence from Human and Monkey Electrophysiology., pp. 687 – 700. MIT Press, 2000.

24. H. H. Jasper, “The ten twenty electrode system of the international federation,” Electroen-cephalography and Clinical Neuroph siology, vol. 10, pp. 371–375, 1958.

25. D. J. Benjamin, J. O. Berger, M. Johannesson, B. A. Nosek, E.-J. Wagenmakers, R. Berk, K. A. Bollen, B. Brembs, L. Brown, C. Camerer, et al., “Redefine statistical significance,” Nature Human Behaviour, vol. 2, p. 4, 2018.

26. V. Amrhein and S. Greenland, “Remove, rather than redefine, statistical significance,” Nature Human Behaviour vol. 2, p.6 – 10, 2018.

27. M. G. Philiastides and S. Gherman, Decision Neuroscience: An Integrative Perspective, ch. Spatiotemporal Characteristics and Modulators of Perceptual Decision-Making, pp. 137 – 148. Academic Press, 2017.

28. R. Oostenveld, P. Fries, E. Maris, and J.-M. Schoffelen, “Fieldtrip: open source software for advanced analysis of MEG, EEG, and invasive electrophysiological data,” Computational intelligence and neuroscience, vol. 2011, p. 1, 2011.

29. S. J. Luck, An introduction to the event-related potential technique. MIT press, 2014.

30. K. J. Friston, D. E. Glaser, R. N. Henson, S. Kiebel, C. Phillips, and J. Ashburner, “Classical and bayesian inference in neuroimaging: applications,” Neurolmage, vol. 16, no. 2, pp. 484–512, 2002.

31. K. J. Friston, W. Penny, C. Phillips, S. Kiebel, G. Hinton, and J. Ashburner, “Classical and Bayesian inference in neuroimaging: theory,” Neurolmage, vol. 16, no. 2, pp. 465–483, 2002.

32. C. Phillips, M. D. Rugg, and K. J. Friston, “Systematic regularization of linear inverse solutions of the eeg source localization problem,” Neurolmage, vol. 17, no. 1, pp. 287–301, 2002.

33. H. R. Heekeren and J. Gold, Neuroeconomics: Decision Making and the Brain, Second Edition, Neural Mechanisms for Perceptual Decision Making, pp. 355–372. Academic Press, 2014.

34. J. A. Maldjian, P. J. Laurienti, R. A. Kraft, and J. H. Burdette, “An automated method for neuroanatomic and cytoarchitectonic atlas-based interrogation of fMRI data sets,” Neurolmage, vol. 19, no. 3, pp. 1233–1239, 2003.

35. K. J. Friston, A. Holmes, J.-B. Poline, C. J. Price, and C. D. Frith, “Detecting activations in PET and fMRI: levels of inference and power,” Neurolmage, vol. 4, no. 3, pp. 223–235, 1996.

## Data Citations

1. Georgie, Y. K., Porcaro, C., Mayhew, S. D., Bagshaw, A. P., and Ostwald, D. Open Science Framework, https://osf.io/q4t8k (2018)

